# A silent Kv channel subunit shapes PV neuron action potential waveform and short-term synaptic plasticity during high-frequency firing

**DOI:** 10.1101/2025.11.06.686832

**Authors:** Sanika Ganesh, Theresa M. Canty, Bernardo L. Sabatini

**Affiliations:** Howard Hughes Medical Institute, Department of Neurobiology, Harvard Medical School, Boston, MA 02115, USA

**Keywords:** PV neuron, potassium channel, silent subunit, GABAergic, inhibition

## Abstract

Fast-spiking parvalbumin-positive (PV) neurons provide precisely timed, context-dependent inhibition within cortical circuits. PV neuron firing properties are specialized among cortical neurons, suggesting that they express a unique complement of ion channels. Here, we identify the PV-specific silent voltage-gated potassium (Kv) channel subunit Kv6.4 (encoded by *Kcng4*), whose role in cortical PV neuron physiology was previously unknown, as a modulator of both intrinsic and synaptic properties. Kv6.4 does not form functional channels on its own but, as shown in prior work, assembles with Kv2 subunits to create heterotetrameric channel complexes, effectively reducing Kv2-mediated delayed rectifier current. We find that *Kcng4* expression is enriched within a distinct *Pvalb*-expressing subclass in primary somatosensory (S1) and motor (M1) cortex and emerges during postnatal development. In PV neurons, Kv6.4 loss reduces action potential (AP) height and width, hyperpolarizes the threshold and interspike potential, and accelerates AP upstroke particularly during repetitive firing. Kv6.4 loss, potentially due to the changes in AP waveform, also alters GABA release and paired-pulse depression at synapses made by PV onto pyramidal (PYR) neurons. The effects of Kv6.4 loss are amplified during high-frequency firing, within the physiological range of fast-spiking PV neurons, likely due to altered repolarization dynamics that accumulate across successive APs. These findings are thus consistent with the function of Kv6.4 in modifying Kv2-mediated delayed rectifier currents. Hence, Kv6.4 tunes the temporal precision of PV inhibitory output, a feature that may be critical for stable excitation-inhibition ratios and adaptive circuit function underlying learning and behavior.

**Significance Statement:** Voltage-gated ion channels are broadly expressed yet serve cell-type-specific functions. We demonstrate that the Kv6.4 silent subunit is selectively expressed by fast-spiking parvalbumin-positive (PV) inhibitory neurons among cortical neurons. In PV neurons, Kv6.4 modulates action potential (AP) waveform and GABA release onto excitatory neurons in a frequency-dependent manner, well within the physiological range of PV firing. This mechanism is likely important for preserving the temporal precision of inhibition, which is particularly critical for fast-spiking interneuron output. By shaping short-term synaptic plasticity during high-frequency firing, Kv6.4 provides a molecular and cell-type-specific mechanism for context-dependent tuning of inhibition within cortical circuits. This function may be especially relevant for sensory processing, experience-dependent plasticity, and the maintenance of excitation-inhibition ratios *in vivo*.

## Introduction

The specific combination of ion channels expressed by each neuron shapes its physiological properties. Large-scale transcriptional analyses have systematically mapped and characterized the molecular signatures of many neuronal classes (1–10), yielding new insights into the variability of ion channel composition across cell classes. These transcriptional data can be integrated with electrophysiological analyses to reveal the mechanisms that govern the intrinsic excitability and synaptic properties of distinct neuron types.

In many brain areas, fast-spiking (FS) interneurons have unique electrophysiological characteristics such as the ability to maintain high rates of action potential (AP) firing. In cerebral cortex, fast-spiking parvalbumin-positive (PV) interneurons deliver precisely timed, potent inhibition by targeting the somatodendritic domain of excitatory pyramidal (PYR) neurons (11). As the largest class of cortical GABAergic neurons, PV neurons preserve network stability by providing cortical gain control (12–14), regulate the temporal precision of PYR neuron firing (15, 16), and gate experience-dependent plasticity (17, 18), especially during critical periods of cortical development (19). Disruptions in PV-cell-mediated inhibition have been implicated in neurological disorders such as epilepsy, schizophrenia, depression, autism spectrum disorder, and Alzheimer’s disease (11, 20).

The diverse functions of PV neurons depend on their capacity for rapid, non-adapting AP firing. Particularly important to sustaining high AP firing rates are potassium (Kv) channels: Kv3 channels support the FS phenotype by facilitating rapid AP repolarization (21–23), whereas Kv1 channels regulate near-threshold excitability by setting AP threshold and first spike latency (24–26). In contrast, the physiological consequences of modulatory potassium channel subunits––nonconducting on their own, but capable of powerfully shaping the behavior of channel complexes––remain largely unexplored (27). These “silent” subunits represent a potentially critical yet understudied mechanism for tuning cellular excitability and circuit dynamics. Notably, their expression is more restricted than that of electrically active channels (28), raising the possibility that their cell-type-specific expression patterns may underlie the functional diversity of neuronal classes.

Multiple single-cell transcriptomic datasets indicate that across cerebral cortical neurons, *Kcng4*, a gene that encodes the nonconducting subunit Kv6.4, is selectively expressed in PV interneurons (3, 10). Prior analyses of cultured and peripheral neurons have demonstrated that Kv6.4 co-assembles with Kv2.1 to generate functional Kv2.1/Kv6.4 channels (29–36). These heterotetrameric channels exhibit a ∼30–40 mV hyperpolarized shift in their voltage dependence of inactivation compared to Kv2.1 homomers, which effectively decreases Kv2 current (29, 30, 32–35). However, given the complex interactions between voltage-sensitive conductances, the impact of Kv6.4 expression on neuronal function is difficult to infer from the biophysical effects of Kv6.4 on Kv2.1 described in heterologous expression systems.

Here, we examine the expression of the Kv6.4 channel subunit in cortical neurons and its contribution to the physiology of cortical PV interneurons. We show that *Kcng4* expression is enriched within *Pvalb*-expressing neurons in primary somatosensory (S1) and motor (M1) cortex. The timing of *Kcng4* expression during postnatal development, together with its spatial distribution, suggest a role in the physiology of mature FS neurons. To examine the function of Kv6.4, we used the Kcng4^Cre^ knock-in mouse line in which Cre recombinase replaces endogenous *Kcng4*, effectively knocking-out the gene. Whole-cell recordings show that Kv6.4 expression modifies AP shape and interspike potentials during repetitive firing without altering AP frequency. At the synaptic level, paired recordings show that a lack of Kv6.4 expression regulates PV inhibition onto nearby PYR neurons during fast spiking, thereby affecting short-term plasticity during rapid firing.

## Results

### Cortical PV neurons selectively express *Kcng4*

The potassium channel Kv6.4 subunit, encoded by *Kcng4*, forms heterotetrameric channels with Kv2.1, encoded by *Kcnb1*. To characterize the expression of these potassium channel subunits in cortical PV neurons, we performed multiplexed fluorescent *in situ* hybridization (FISH) to examine the expression of *Kcng4*, *Kcnb1*, and *Pvalb* (n = 6 mice with 18 sections and 36 hemispheres per region, 2–4 sections per mouse) in M1 and S1 in wildtype C57BL/6J mice (**Fig. 1 *A–B***).

**Fig. 1.**
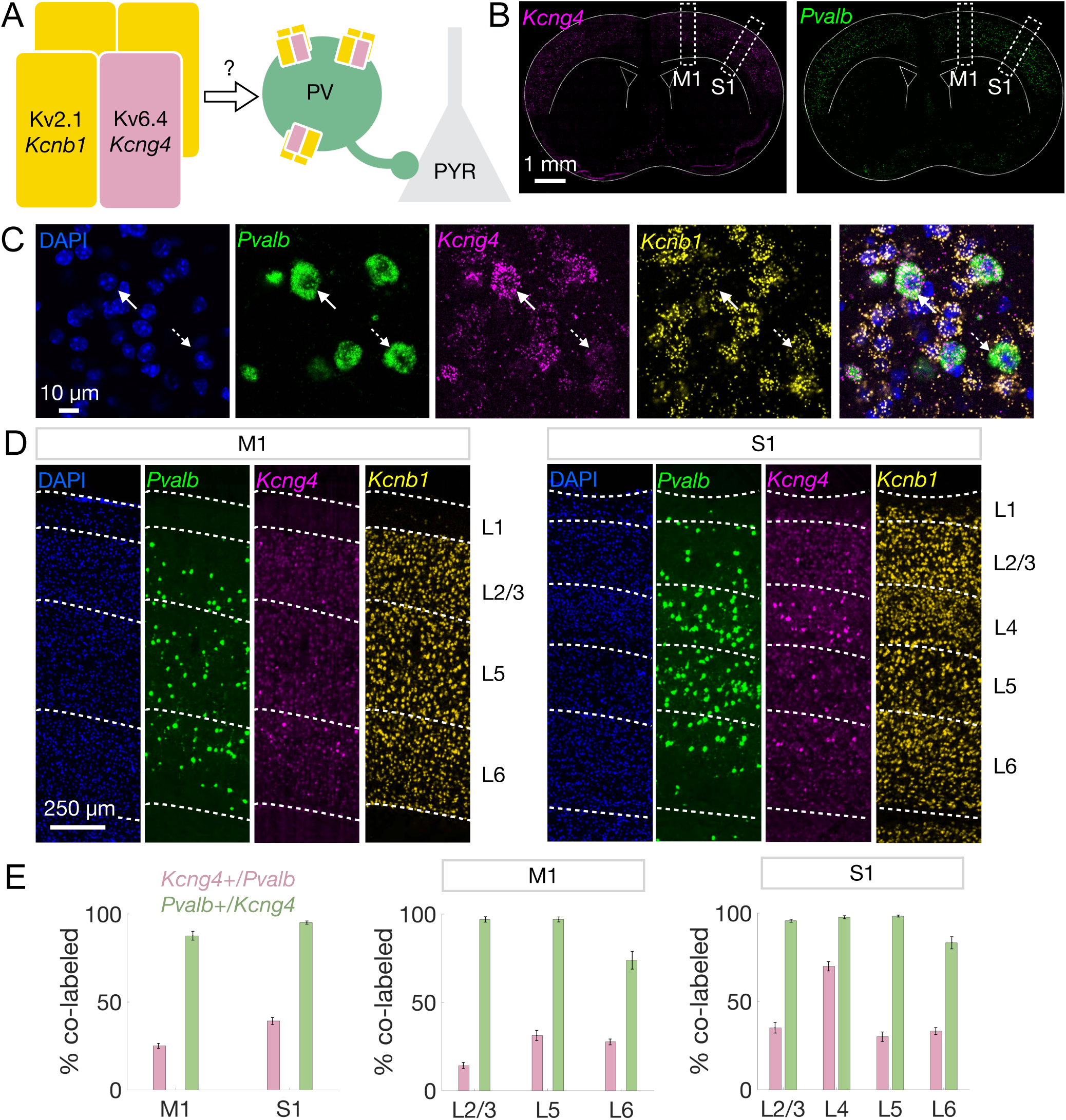
Cortical PV neurons selectively express *Kcng4*. **A,** Schematic illustrating the hypothesis that Kv6.4 (encoded by *Kcng4*) confers cell-type-specific function by forming heterotetrameric channels with Kv2.1 (encoded by *Kcnb1*). We tested whether Kv6.4 modulates PV neuron excitability and PV inhibition of excitatory pyramidal (PYR) neurons. **B**, Coronal brain sections displaying the spatial distribution of *Kcng4* (magenta, *left*) and *Pvalb* (green, *right*) mRNA across cortex detected by FISH. Subsequent analyses focus on primary motor (M1) and somatosensory (S1) cortical regions, indicated by white dashed boxes. **C**, Representative images of individual cortical cells labeled with DAPI (blue) and FISH against *Pvalb* (green), *Kcng4* (magenta), and *Kcnb1* (yellow), followed by a merged view (*left* to *right*). Solid white arrows indicate a cell that co-expresses *Pvalb, Kcnb1,* and *Kcng4*, and dashed white arrows indicate a cell that co-expresses *Pvalb* and *Kcnb1*. **D**, Representative images of M1 and S1 showing the spatial distribution of *Kcng4*, *Pvalb*, and *Kcnb1* mRNA. White dashed lines and text labels denote individual cortical layers (L). **E**, Quantification of co-labeled cells displaying the percentage of *Pvalb*-expressing cells that are *Kcng4*+ (pink) and the percentage of *Kcng4*-expressing cells that are *Pvalb*+ (green) in M1 and S1 (n = 6 mice with 18 sections and 36 hemispheres per region, 2–4 sections per mouse) across all layers (*left*). Quantification by cortical layer in M1 (*center*) and S1 (*right*). Mean ± SEM displayed.

We detected expression of *Kcng4* across cortical layers in M1 and S1, roughly mirroring that of *Pvalb* (**Fig. 1 *C–D***). High-resolution images demonstrate *Kcng4* and *Pvalb* co-expression within individual cells (**Fig. 1*C***). Indeed, nearly all *Kcng4*+ cells express *Pvalb* (M1: 0.88 ± 0.025, S1: 0.95 ± 0.009), and a substantial proportion of *Pvalb*+ neurons express *Kcng4* (M1: 0.25 ± 0.014, S1: 0.39 ± 0.020), with higher co-expression of *Pvalb* and *Kcng4* in S1 compared to M1 (**Fig. 1*E*** and **Dataset S1**). These cortical co-expression patterns are reminiscent of those observed in PV neurons of the external globus pallidus (GPe), where ∼90% of *Kcng4*+ neurons express *Pvalb*, and ∼30% of *Pvalb*+ neurons express *Kcng4* (37). In both M1 and S1, co-expression patterns were consistent across layers (L; **Fig. 1*D***). However, we noted high enrichment of *Kcng4* in S1 L4 *Pvalb*+ neurons (S1 L4: 0.70 ± 0.027; **Fig. 1*E*** and **Dataset S1**). We found that almost every neuron that co-expressed *Kcng4* and *Pvalb* also expressed *Kcnb1* (*Kcnb1*: 0.99 ± 0.003; **Dataset S1**). While most functional studies of Kv6.4 have reported its assembly with Kv2.1 (29–35, 38), Kv2.1 and Kv2.2 (encoded by *Kcnb2*) can be expressed independently (39) and interact with Kv6.4 (36). Both *Kcnb1* and *Kcnb2* are expressed by PV neurons (**Fig. S1*C***).

The results of these FISH analyses are consistent with spatial transcriptomic data from the Allen Brain Cell (ABC) Atlas (10), which confirm that *Kcng4* is selectively expressed in *Pvalb*+ neurons in M1 and S1 (**Fig. S1*A***). We also noted that *Kcng4*+ neurons comprise a unique transcriptionally defined supertype (supertype 207) of *Pvalb*+ interneurons (**Fig. S1*B***), as defined by the ABC Atlas (10).

### Spatiotemporal patterns of Kcng4^Cre^ activation

*Pvalb* expression emerges around postnatal day (P) 12, and PV neuron maturation restricts critical periods of heightened sensory-motor cortical plasticity (18, 19, 25, 40). Therefore, we examined the onset of Kv6.4 expression. We generated Kcng4^Cre^; Ai14 mice by crossing a transgenic mouse line in which Cre is knocked into the *Kcng4* locus (Kcng4^Cre^), resulting in a functional knock-out, to a reporter line that expresses tdTomato in a Cre-dependent manner (Ai14). We used tdTomato fluorescence to compare Kcng4^Cre^ activation in M1 and S1 of young (2 weeks) and adult (5+ weeks) heterozygous (Kcng4^Cre/WT^; Ai14) and homozygous (Kcng4^Cre/Cre^; Ai14) mice (**Fig. 2*A***). Because Cre replaces the endogenous *Kcng4* coding sequence in Kcng4^Cre^ mice, heterozygous mice retain one functional *Kcng4* allele, and homozygous mice lack functional *Kcng4* alleles entirely. We therefore expected Cre-driven tdTomato expression to exhibit a gene-dosage effect, with higher Cre reporter activation in homozygous than heterozygous mice. Consistent with this expectation, tdTomato fluorescence was modestly higher in homozygous mice (**Fig. 2 *B–D*).** Importantly, both genotypes showed weak labeling in young mice and strong labeling in adult mice in M1 and S1 (**Fig. 2 *B–D*).** High-resolution imaging at the 2-week timepoint revealed that tdTomato-labeled cortical cells in young animals exhibit astrocytic morphology, characterized by extensive arborization and a lack of polarized processes (**Fig. S2 *A–C***). These results suggest that there is minimal Kcng4^Cre^ activation in PV neurons at early developmental stages, and that Kv6.4 expression arises after maturation of PV neurons, specifically in M1 and S1.

**Fig. 2.**
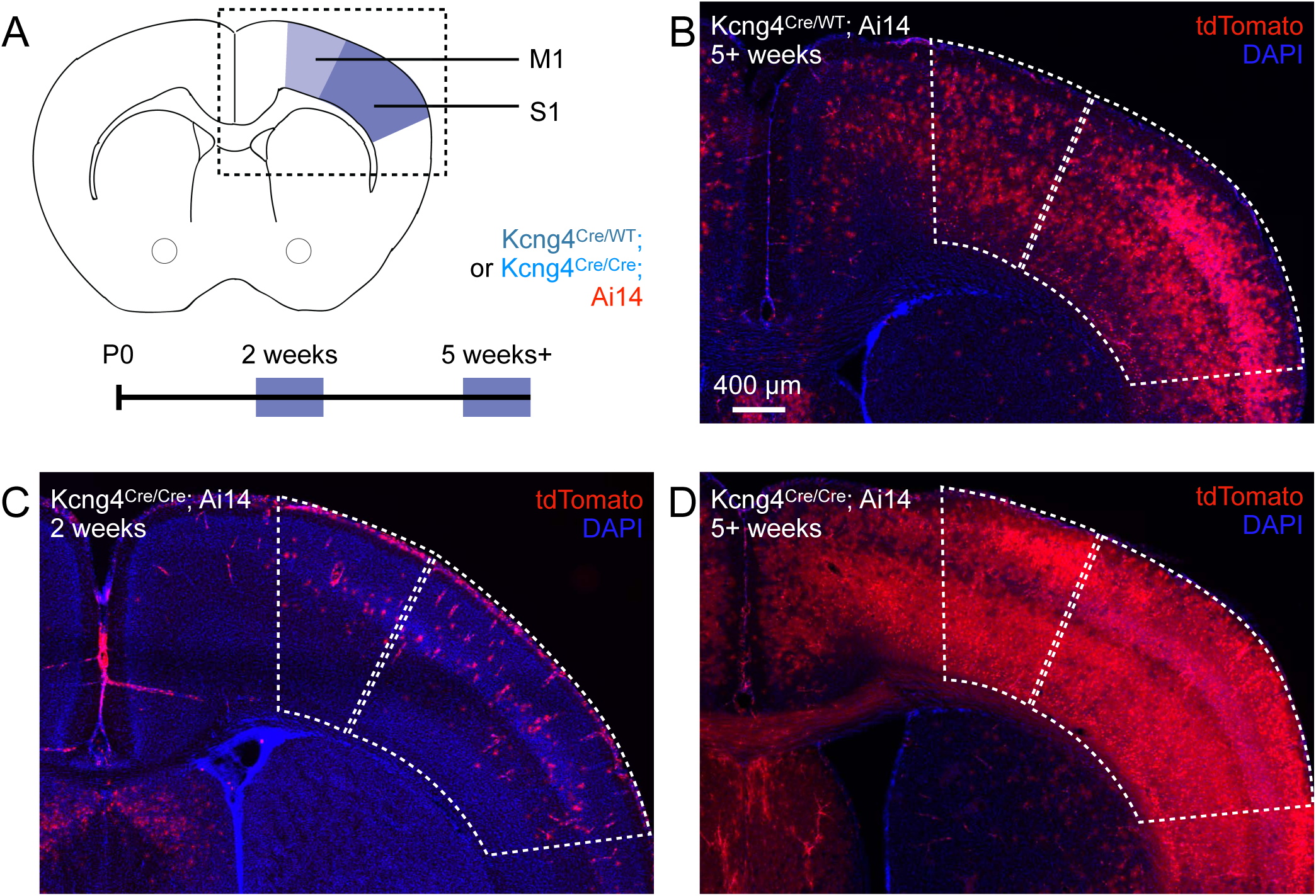
Age dependence of Kcng4^Cre^ activation. **A**, Schematic of the brain regions analyzed to assess Kcng4^Cre^ activation at juvenile (2 weeks) and adult (5+ weeks) timepoints in heterozygous (Kcng4^Cre/WT^; Ai14) and homozygous (Kcng4^Cre/Cre^; Ai14) mice. Analyses focused on primary motor (M1) and somatosensory (S1) cortex, as indicated. **B–D**, Representative images showing tdTomato fluorescence in M1 and S1, as indicated in panel A, for Kcng4^Cre/WT^; Ai14 mice at 5+ weeks **(B)**, and Kcng4^Cre/Cre^; Ai14 mice at 2 weeks **(C)** and 5+ weeks **(D)**.

In adult mice, Kcng4^Cre^-mediated expression of tdTomato occurs in brain regions that contain fast-spiking neurons, such as the cerebral cortex, GPe, and cerebellum (**Fig. S3**). Indeed, *Kcng4*+ neurons have previously been identified as a subclass of PV neurons in the GPe (37). In the cerebellum, *Kcng4* expression is enriched in fast-spiking Purkinje cells (**Fig. S1*D***) as may be expected from our observations, given that PV neurons include some but not all Purkinje cells (6). The spatial distribution of tdTomato+ neurons is thus consistent with Kv6.4 expression in fast-spiking neuronal populations across multiple brain regions. Together, the spatial and temporal patterns of Kcng4^Cre^ activation raise the possibility that Kv6.4 shapes the fast-spiking phenotype of PV neurons.

### Kv6.4 modulates AP height, AP half-width, and interspike potential in PV neurons

Given that Kv6.4 reduces delayed rectifier Kv2-mediated current in heterologous expression systems (29, 30, 32–35), the removal of Kv6.4 in PV neurons is expected to increase net Kv2 current. This could have widespread effects on the intrinsic properties of PV neurons, such as AP shape and interspike potential. Thus, to study the impact of Kv6.4 on PV neuron physiology, we performed whole-cell current-clamp recordings in fast-spiking neurons from L2/3 wildtype *Kcng4* (n = 18 cells in 4 mice, PV^Cre/WT^; Ai14), heterozygous *Kcng4* knock-out (n = 16 cells in 4 mice, Kcng4^Cre/WT^; Ai14), and homozygous *Kcng4* knock-out (n = 20 cells in 4 mice, Kcng4^Cre/Cre^; Ai14) mice carrying two (WT/WT), one (Cre/WT), or no (Cre/Cre) copies of *Kcng4*, respectively. In each genotype, recordings were made from tdTomato-expressing neurons in age-matched adult animals. Because S1 *Pvalb+* neurons are more likely to express *Kcng4* than M1 *Pvalb+* neurons (**Fig. 1*E***), we conducted electrophysiological recordings in S1. We used a series of current injections to measure voltage (–200 to 200 pA, 40 pA increments) and AP (200 to 1200 pA, 100 pA increments) responses across all three genotypes, blinded to heterozygous and homozygous animals from which acute brain slices were prepared (**Fig. 3*A***). We used littermates to compare heterozygous and homozygous animals. Although we observed no significant gross differences in current–voltage or current–firing rate relationships across genotypes (**Fig. 3*B*** and **Dataset S2**), we observed several significant differences in intrinsic properties between wildtype, heterozygous, and homozygous mice, in line with our expectation that Kv6.4 loss alters Kv2-mediated delayed rectifier current.

**Fig. 3.**
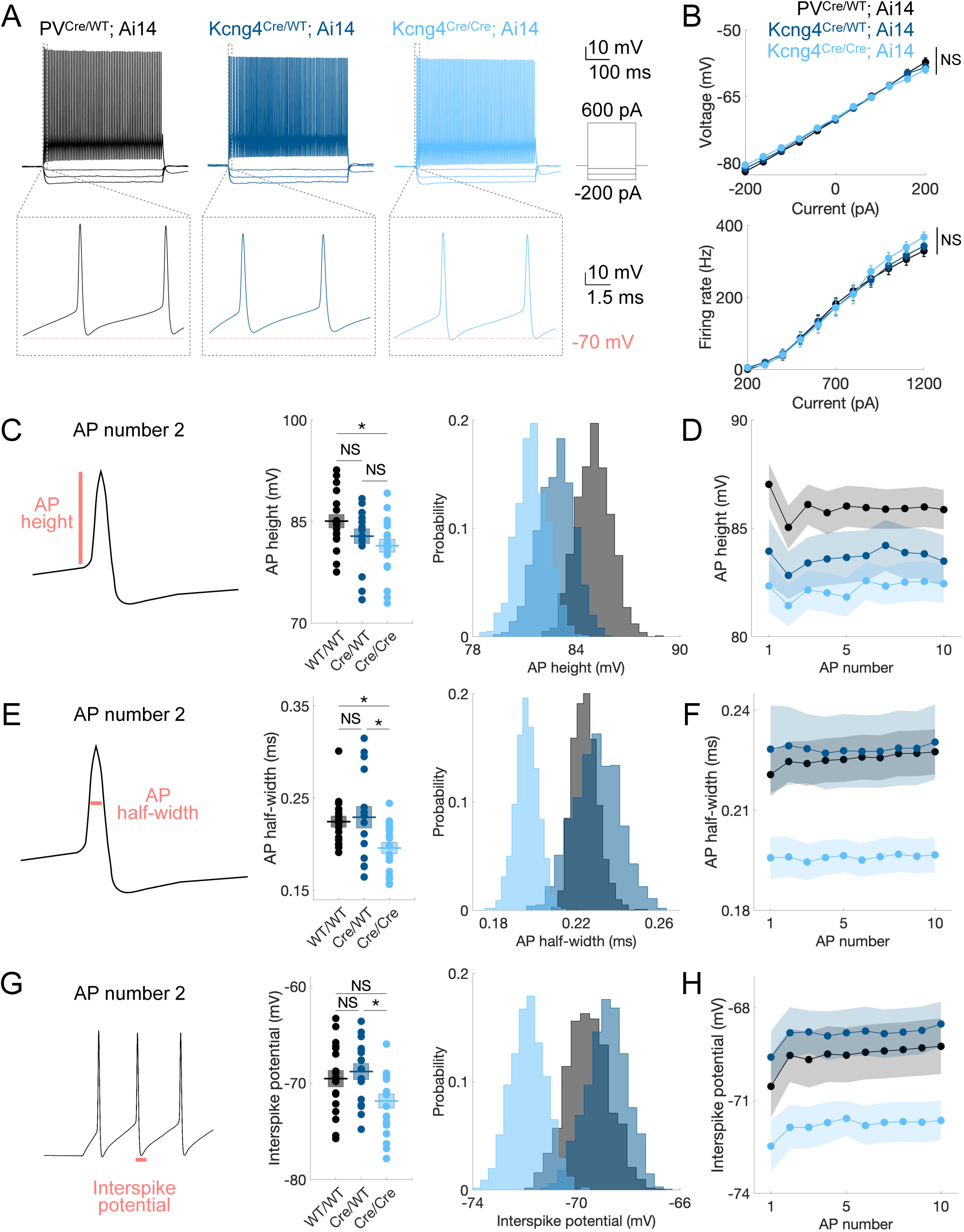
Kv6.4 modulates AP height, AP half-width, and interspike potential in PV neurons. **A**, Representative whole-cell current-clamp recordings from L2/3 fast-spiking neurons in S1 of mice with two (PV^Cre/WT^; Ai14), one (Kcng4^Cre/WT^; Ai14), and no (Kcng4^Cre/Cre^; Ai14) active alleles of *Kcng4*. Voltage responses to a series of current injections (–200, –120, –40, and 600 pA) are shown in the top row. An expanded view of the first two action potentials in response to a 600 pA current injection is shown in the bottom row, aligned to a membrane potential of –70 mV. **B**, Voltage (mV, *top*) and firing rate (Hz, *bottom*) responses to stepwise current injections for wildtype (n = 18 cells in 4 mice), heterozygous (n = 16 cells in 4 mice), and homozygous (n = 20 cells in 4 mice) groups. **C**, Schematic of action potential (AP) height (*left*) and its group mean measured for the 2^nd^ AP (mV, *center*) in the first current step that elicited at least 10 APs. Bootstrapped distribution of the means (*right*). **D**, Mean values across the first 10 APs (mV) in a response that elicited at least 10 APs. **E–H**, As in **C** and **D** for AP half-width (ms) (**E-F**) and interspike potential (mV) (**G-H**). Mean ± SEM displayed. * *P* < 0.05; NS: not statistically significant. Repeated-measures ANOVA (Greenhouse-Geisser corrected) or one-way ANOVA (with Tukey-Kramer multiple comparisons). Bootstrapping (1,000 iterations) was used to visualize sample mean distributions.

Indeed, AP height, AP half-width, and interspike potential differed significantly between groups (**Fig. 3 *C–F*** and **Dataset S2)**. To standardize analysis across cells with varying rheobase, we analyzed the first trace containing at least 10 APs and restricted analysis to the 2^nd^ and 10^th^ APs in these traces to survey early and later phases of the AP train. AP height was lower in homozygous mice compared to wildtype for the 2^nd^ AP (WT/WT = 85.05 ± 1.00 mV, Cre/WT = 82.84 ± 1.10 mV, Cre/Cre = 81.44 ± 0.95 mV; one-way ANOVA, *p* = 0.041; Tukey–Kramer post hoc comparisons: WT/WT vs Cre/WT *p* = 0.297, WT/WT vs Cre/Cre *p* = 0.032, Cre/WT vs Cre/Cre *p* = 0.597), and for the 10^th^ AP (**Fig. 3 *C–D*** and **Dataset S2**). Half-width was also significantly reduced in homozygous mice, both for the 2^nd^ AP (WT/WT = 0.22 ± 0.01 ms, Cre/WT = 0.23 ± 0.01 ms, Cre/Cre = 0.20 ± 0.01 ms; one-way ANOVA, *p* = 0.007; Tukey–Kramer post hoc comparisons: WT/WT vs Cre/WT *p* = 0.911, WT/WT vs Cre/Cre *p* = 0.030, Cre/WT vs Cre/Cre *p* = 0.013) and the 10^th^ AP (**Fig. 3 *E–F*** and **Dataset S2**). Additionally, the interspike potential, defined as the most negative potential between APs, was more negative in homozygous mice for the 2^nd^ AP (WT/WT = –69.55 ± 0.86 mV, Cre/WT = –68.81 ± 0.83 mV, Cre/Cre = –71.87 ± 0.72 mV; one-way ANOVA, *p* = 0.022; Tukey–Kramer post hoc comparisons: WT/WT vs Cre/WT *p* = 0.805, WT/WT vs Cre/Cre *p* = 0.098, Cre/WT vs Cre/Cre *p* = 0.026) and the 10^th^ AP (**Fig. 3 *G–H*** and **Dataset S2**). Bootstrapping analysis further confirmed differences in the distributions across groups (**Fig. 3 *C, E,* and *G***). These results indicate that Kv6.4 loss decreases AP amplitude and duration and deepens the interspike potential in PV neurons. These effects are consistent with the known mechanism through which Kv6.4 reduces Kv2 current, namely by shifting heterotetramer inactivation to more negative potentials.

Additionally, we examined AP threshold, AP maximum slope, afterhyperpolarization (AHP) amplitude for a train of APs, and sag voltage across the three groups. We found that the three groups differed significantly in AP threshold for the 10^th^ AP (WT/WT = –42.14 ± 0.81 mV, Cre/WT = –42.00 ± 0.58 mV, Cre/Cre = –44.14 ± 0.60 mV; one-way ANOVA, *p* = 0.046; Tukey–Kramer post hoc comparisons: WT/WT vs Cre/WT *p* = 0.988, WT/WT vs Cre/Cre *p* = 0.090, Cre/WT vs Cre/Cre *p* = 0.076), but not for the 2^nd^ AP (**Fig. S4 *A–B*** and **Dataset S2**), suggesting that Kv6.4 loss contributes to maintaining a lower AP threshold particularly over the course of repetitive firing (**Fig. S4*B***). AP maximum slope was also significantly higher at the 10^th^ AP in homozygous mice (WT/WT = 452.02 ± 11.98 V/s, Cre/WT = 432.02 ± 22.33 V/s, Cre/Cre = 488.23 ± 13.40 V/s; one-way ANOVA, *p* = 0.047; Tukey–Kramer post hoc comparisons: WT/WT vs Cre/WT *p* = 0.666, WT/WT vs Cre/Cre *p* = 0.233, Cre/WT vs Cre/Cre *p* = 0.042), but no differences were observed at the 2^nd^ AP (**Fig. S4 *C–D*** and **Dataset S2**). Thus, when Kv6.4 is absent, fast-spiking neurons exhibit an increased upstroke velocity during repetitive firing. There were no significant differences in the AHP amplitude (**Fig. S4 *E–F*** and **Dataset S2**) or sag voltage (**Fig. S4 *G–H*** and **Dataset S2**), which is consistent with the limited role of Kv6.4 in the conductances underlying these parameters. As discussed previously, Kv6.4 loss is predicted to increase net Kv2-mediated repolarizing drive, and consistent with this prediction, the interspike potential is hyperpolarized in PV neurons that do not express *Kcng4* (**Fig. 3 *G-H*** and **Dataset S2**). This deeper hyperpolarization is expected to enhance the recovery of voltage-gated sodium (Nav) channels from inactivation during repetitive firing (41). Thus, in the absence of Kv6.4, enhanced Nav channel availability likely contributes to the more hyperpolarized AP threshold and faster AP upstroke velocity that we observe during sustained firing in PV neurons.

To determine whether the differences between groups generalize across firing rates, we analyzed AP height, half-width, interspike potential, threshold potential, and maximum slope across all recordings, with data organized into bins containing comparable numbers of APs (**Fig. S5*A***). We found that differences in genotype persist across firing rates (**Fig. S5 *B–F***), indicating that these effects are not limited to a single firing rate regime. Thus, group differences in these metrics are robust to the trace selection criterion described previously (2^nd^ and 10^th^ AP within the first trace containing ≥10 APs for the analysis shown in **Fig. 3** and **Fig. S4**). The shorter AP duration in homozygous animals is not attributable to adaptation of the interspike interval (ISI; ms), as the groups exhibit a similar change (Δ) in ISI between early and late spikes (ISI49-50 – ISI2-3) in recordings containing ≥50 APs (**Fig. S5*G***). Notably, homozygous animals display attenuated within-trace adaptation in AP half-width and interspike potential between early and late spikes (AP50 – AP2; **Fig. S5 *H–I***). This reduced adaptation in AP half-width becomes more pronounced at higher firing rates (**Fig. S5*H***). Hence, Kv6.4 loss deepens the interspike potential and shortens action potentials across all firing rates examined, with increasing effects at higher firing frequencies.

Because the S1 L2/3 WT PV population includes both *Kcng4*-expressing and non-expressing neurons, differences between wildtype, heterozygous, and homozygous animals may reflect differences in PV neuron subtypes, rather than specific physiological effects of removing the Kv6.4 silent subunit. Specifically, roughly 35% of S1 L2/3 PV neurons express *Kcng4*. However, phenotypes that differ between the heterozygous and homozygous mice isolate effects within neurons that typically express *Kcng4*. These include AP half-width, interspike potential, and AP maximum slope, indicating that these alterations in AP waveform reflect Kv6.4 loss rather than variability between PV subclasses or differences in mouse genetic background. In contrast, differences in AP height and AP threshold may be influenced by the inclusion of *Kcng4* non-expressing neurons in the WT group. Nonetheless, these effects are consistent with the expected mechanisms through which Kv6.4 loss modifies Kv2 repolarization and associated Nav availability, suggesting that these effects arise from physiological changes rather than PV neuron subtype variability.

Hence, especially during repetitive firing, Kv6.4 deletion yields more negative interspike and threshold potentials, faster AP upstrokes, and narrower and smaller APs. These altered intrinsic properties are consistent with the expected increase in Kv2 repolarizing current and enhanced recovery of Nav current. Notably, these alterations do not result in a higher firing frequency but rather appear to refine the response of PV neurons during periods of high activity. Therefore, Kv6.4 shapes the AP waveform in PV neurons, particularly during high-frequency firing.

### Kv6.4 regulates paired-pulse depression at PV–PYR synapses during high-frequency AP firing

Motivated by the possibility that Kv6.4-dependent changes in PV action potential waveform could alter GABA release, we next investigated inhibitory transmission at PV–PYR synapses. To study the consequences of Kv6.4 ablation on inhibitory synapses, we performed paired whole-cell recordings in L2/3 PV neurons and connected PYR neurons (**Fig. 4*A***). This analysis included wildtype (n = 11 pairs in 9 mice) and homozygous (n = 10 pairs in 8 mice) groups. We evoked action potentials in PV neurons by current injection under current-clamp and simultaneously measured the resulting inhibitory postsynaptic currents (IPSCs) in neighboring PYR neurons under voltage-clamp (**Fig. 4*B***). Excitatory currents were pharmacologically blocked. PYR neuron recordings used a high-chloride cesium-based internal solution to amplify inhibitory currents, resulting in inward GABAergic currents at resting membrane potentials. Synaptic currents were also converted to conductance to isolate postsynaptic channel activation independent of recording conditions (see ***Synaptic Properties*** in ***Materials and Methods***). We did not identify differences in IPSC peak amplitude, whether assessed as current or conductance, or in the passive properties (membrane resistance, membrane capacitance, or membrane time constant) of PYR neurons between groups (**Fig. 4*C*** and **Dataset S3**).

**Fig. 4.**
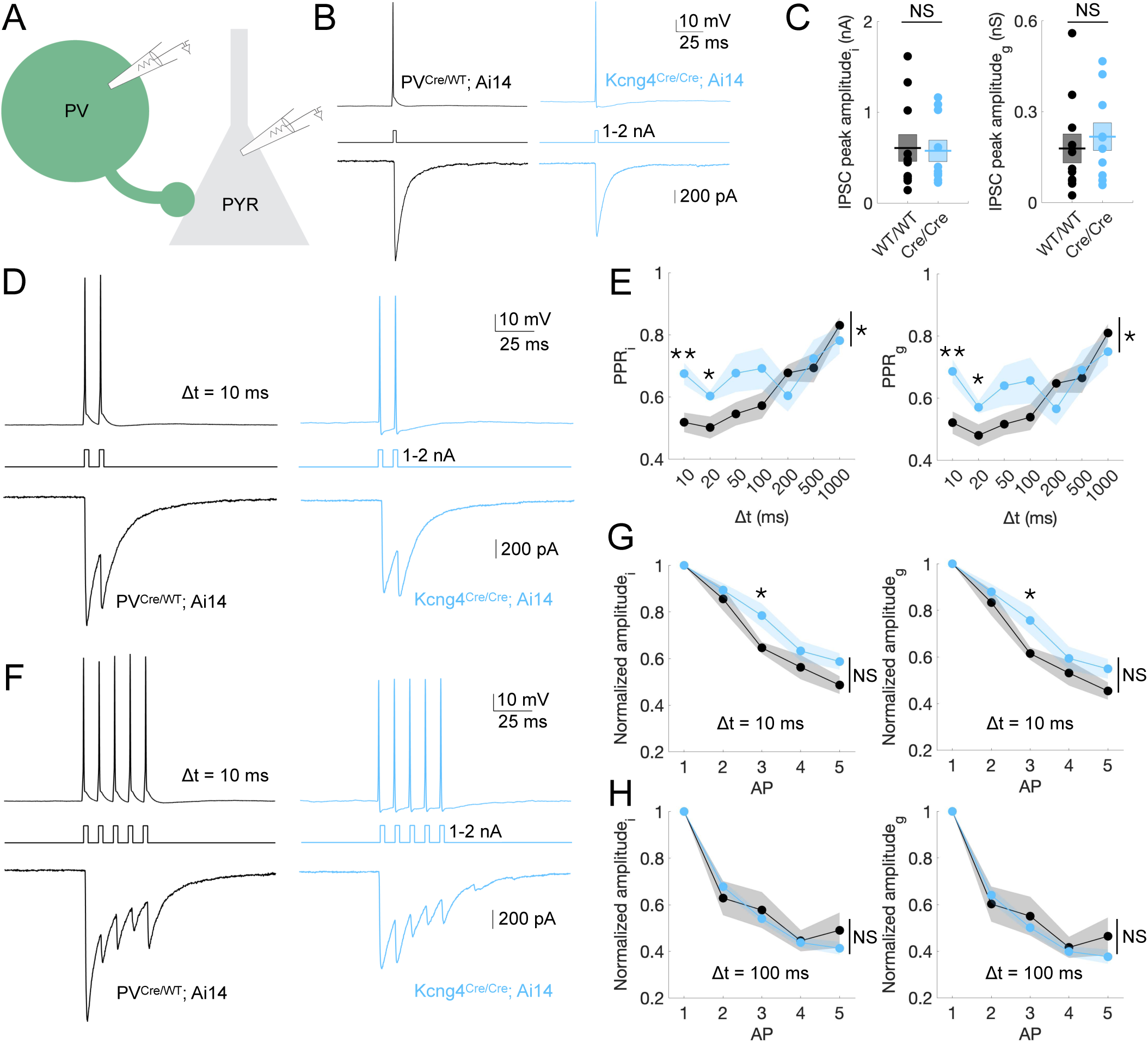
Kv6.4 regulates paired-pulse depression at PV-PYR synapses during high-frequency AP firing. **A,** Schematic of paired whole-cell recordings from PV and PYR neurons. **B,** Representative whole-cell voltage-clamp recordings from L2/3 PYR neurons in S1 of mice with two (PV^Cre/WT^; Ai14) and no (Kcng4^Cre/Cre^; Ai14) active alleles of *Kcng4*. APs were evoked in tdTomato-positive, fast-spiking neurons by brief current injections (1–2 nA, 1 ms), resulting in inhibitory postsynaptic currents (IPSCs) in synaptically connected PYR neurons. A representative response to a single AP is displayed for each group. **C,** IPSC peak amplitude measured as current (*i*; *left,* nA) or converted to conductance (*g*; *right,* nS) for wildtype (n = 11 pairs in 9 mice) and homozygous (n = 10 pairs in 8 mice) groups. **D**, Representative response to two presynaptic action potentials (paired-pulse stimulation) delivered with an interstimulus interval of 10 ms, recorded as in **B**. **E**, Paired-pulse ratio (PPR) of inhibitory current (*i*; *left*) and conductance (*g*; *right*) across groups for ISIs of 10, 20, 50, 100, 200, 500, and 1000 ms. **F**, Representative response to five presynaptic action potentials delivered with an interstimulus interval of 10 ms, recorded as in **B** and **D**. **G**, Amplitude of inhibitory current (*i*; *left*) and conductance (*g*; *right*) across groups, normalized to response from the first AP, for an ISI (Δt) of 10 ms. **H**, As in **G** for an ISI of 100 ms. Mean ± SEM displayed. ** *P* < 0.005, * *P* < 0.05; NS: not statistically significant. Unpaired two-tailed t-tests or repeated-measures ANOVA (Greenhouse-Geisser corrected).

We found that Kv6.4 modulates synaptic depression during high-frequency firing, as indicated by altered inhibitory currents in response to paired-pulse stimulation. Specifically, we recorded IPSCs in PYR neurons triggered by pairs of action potentials in PV neurons with varying interstimulus intervals (Δt = 10, 20, 50, 100, 200, 500, 1000 ms; **Fig. 4*D***). The amplitude of overlapping IPSCs was quantified by subtracting the residual decay of the first IPSC from the peak of the second (see ***Synaptic Properties*** in ***Materials and Methods***). We observed that the paired-pulse ratio of inhibition (PPRi) and conductance (PPRg) is significantly higher in mice that lack Kv6.4 compared to control mice, but only at short interstimulus intervals, Δt = 10 ms (PPRi: WT/WT = 0.52 ± 0.031, Cre/Cre = 0.68 ± 0.035; unpaired two-tailed t-test, *p* = 0.003; PPRg: WT/WT = 0.52 ± 0.036, Cre/Cre = 0.69 ± 0.038; unpaired two-tailed t-test, *p* = 0.005) and Δt = 20 ms (PPRi: WT/WT = 0.50 ± 0.036, Cre/Cre = 0.60 ± 0.016; unpaired two-tailed t-test, *p* = 0.022; PPRg: WT/WT = 0.48 ± 0.035, Cre/Cre = 0.57 ± 0.020; unpaired two-tailed t-test, *p* = 0.043) (**Fig. 4*E*** and **Dataset S3**). This indicates that removal of Kv6.4 alters presynaptic GABA release properties of PV neurons. The observed differences are consistent with a reduction in probability of GABA release per release site in mice lacking *Kcng4*, although paired-pulse measurements alone cannot distinguish reduced release probability from alternative and not mutually exclusive mechanisms such as enhanced short-term facilitation. In either case, differences in presynaptic properties between groups, such as Ca^2+^ entry triggering neurotransmitter release, can be explained by the altered spike waveform observed in knockout neurons (**Fig. 3**, **Fig. S5**). Moreover, the relationship between paired-pulse ratio and interstimulus interval was significantly different between genotypes (repeated-measures ANOVA; current, *p* = 0.032; conductance, *p* = 0.038; **Dataset S3**), indicating that synaptic depression varied across interstimulus intervals as a function of Kv6.4 expression. Thus, paired-pulse depression is reduced at PV–PYR synapses in the absence of Kv6.4 expression for presynaptic APs firing faster than ∼50 Hz, which is well below the maximal firing rates (hundreds of Hz) of cortical PV neurons *in vivo* (42).

We observed similar trends in the amplitude of inhibitory currents elicited by a 5-pulse train, normalized to the first pulse: effects were only apparent at the shortest interstimulus interval (**Fig. 4 *F–H*** and **Dataset S3**). As the decay of preceding IPSCs was not corrected in the analysis of 5-pulse train measurements (see ***Synaptic Properties*** in ***Materials and Methods***), the differences between groups in this analysis may appear less pronounced than in the paired-pulse analysis. At Δt = 10 ms, both current (WT/WT = 0.65 ± 0.025, Cre/Cre = 0.78 ± 0.051; unpaired two-tailed t-test, *p* = 0.022) and conductance (WT/WT = 0.61 ± 0.027, Cre/Cre = 0.76 ± 0.059; unpaired two-tailed t-test, *p* = 0.035) were significantly higher for the third pulse in homozygous mice. No differences were detected for other pulses or interstimulus intervals in individual comparisons, and repeated-measures ANOVA across all five pulses revealed no significant overall effect of genotype, although a trend toward higher normalized amplitudes in the absence of Kv6.4 was observed at the shortest interstimulus interval (**Fig. 4 *G–H***, **Fig. S*6***, and **Dataset S3**).

As with many knockout-based approaches, interpretation of genotype-dependent effects must acknowledge that the targeted gene is not uniformly expressed across all neurons of the targeted cell type. As discussed previously, WT recordings therefore include a mix of *Kcng4*-expressing and non-expressing neurons and may sample from different PV subpopulations. Nevertheless, the differences in repolarization observed between heterozygous and homozygous neurons (**Fig. 3**, **Fig. S5**), both of which normally express *Kcng4* and therefore control for PV subtype diversity, offer a plausible mechanistic basis for the synaptic phenotype. Specifically, the shortening of action potentials caused by Kv6.4 loss is expected to alter presynaptic calcium entry in ways that could influence GABA release.

Together, these results demonstrate that Kv6.4 selectively tunes synaptic depression at PV–PYR synapses during high-frequency activity, without influencing steady-state inhibition or IPSCs at longer interstimulus intervals. These data reveal a function for Kv6.4 in regulating the precision of inhibitory signaling at cortical PV–PYR synapses under fast-spiking conditions.

## Discussion

Our findings reveal that Kv6.4 expression shapes both the intrinsic properties and synaptic output of PV neurons during high-frequency firing (**Fig. 5**). Altered repolarization dynamics likely emerge at the interspike potential and accumulate during repetitive firing, progressively modifying the AP waveform, reducing its duration, amplitude, and threshold while increasing upstroke velocity. These waveform changes likely modify synaptic release during high-frequency firing, consistent with our observation of altered paired-pulse ratios at short interstimulus intervals. Thus, Kv6.4 regulates PV inhibition onto pyramidal neurons in a frequency-dependent manner.

**Figure 5.**
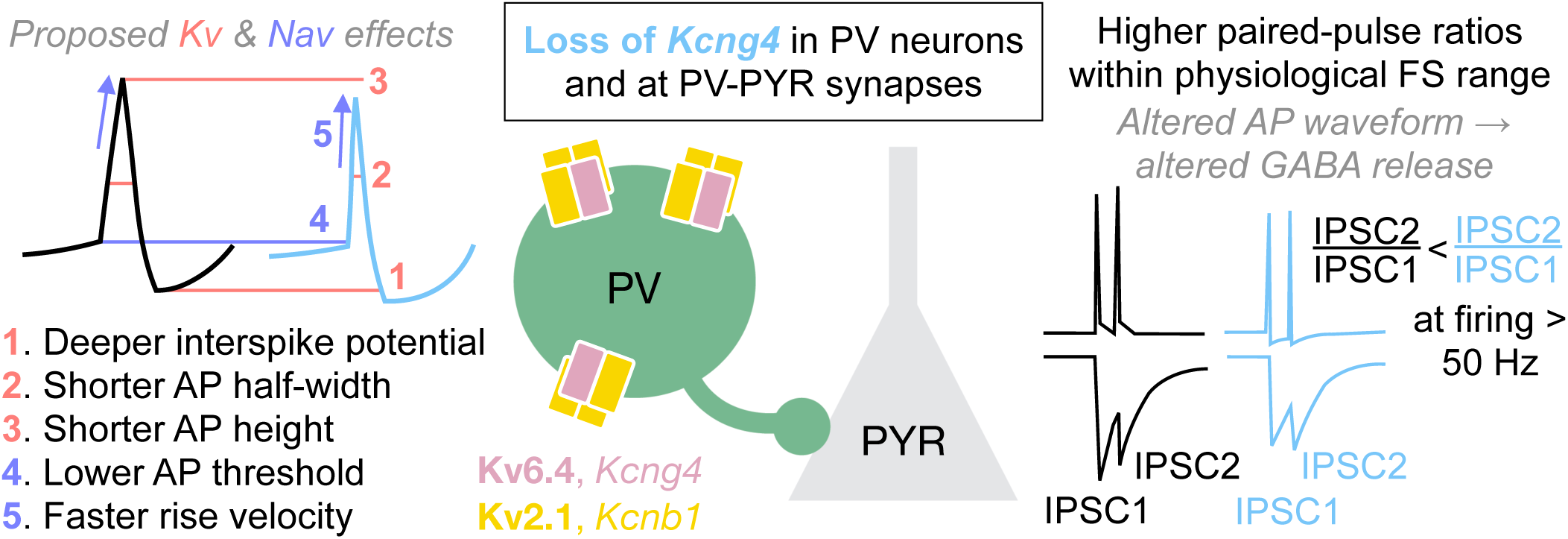
Model: Kv6.4 alters PV neuron waveform and inhibition of PYR neurons. Kv6.4 (*Kcng4*) forms heterotetrameric potassium channel complexes in PV neurons. The absence of Kv6.4 results in (1) deeper interspike potentials, (2) shorter AP half-width, and (3) shorter AP height, likely due to its effects on Kv2 currents. Loss of Kv6.4 also results in (4) lower AP threshold and (5) faster rise velocity, which we propose is due to a resulting increase in Nav channel availability. We propose that these waveform changes lead to an increase in paired-pulse ratios of inhibition evoked in postsynaptic PYR neurons at short interstimulus intervals. Thus, Kv6.4 regulates presynaptic GABA release during high-frequency firing and thereby regulates the temporal precision of PV inhibitory output.

The effects of Kv6.4 are likely mediated through its interactions with Kv2 subunits. The altered interspike potential observed in Kv6.4 knock-out neurons is consistent with prior findings that Kv6.4 subunits influence the inactivation properties of Kv2 heterotetramers (29, 30, 32–35). Kv2 current provides repolarizing drive that accumulates during high-frequency trains due to its slow kinetics, preventing AP broadening (28, 43–45). Loss of Kv2.1 prolongs APs and broadens Ca^2+^ transients specifically during high-frequency firing (43). In line with this, Kv6.4 likely broadens APs in PV neurons by reducing Kv2 current density, as it promotes the inactivation of its Kv2 partners during repolarization (29, 30). The contribution of Kv6.4 likely accompanies secondary effects to other potassium and sodium channels that further influence the spike train (41). Kv6.4 depolarizes the interspike potential and broadens the AP waveform in PV neurons, which we propose may subsequently elevate the initial probability of GABA release at PV–PYR cell synapses, resulting in the enhanced paired-pulse depression observed (**Fig. 5**). Subtle alterations in AP waveform, and consequently in presynaptic Ca^2+^ entry, are known to exert supralinear effects on postsynaptic current amplitude (46–48), underscoring the sensitivity of neurotransmitter release to AP narrowing or broadening. In PV neurons, Ca^2+^ channels and GABA release are tightly coupled to rapidly translate changes in AP waveform into graded inhibition (11). Deletion of both *Kcng4* alleles results in a ∼10% change in AP width (**Fig. 3** and **Dataset S2**) but a ∼30% change in paired-pulse ratio (**Fig. 4** and **Dataset S3**). This ∼3-fold amplification of Kv6.4-dependent effects is expected if AP broadening is also present in the presynaptic boutons of PV neurons. Kv3 channels have long been recognized as the primary regulators of rapid firing in PV neurons and synaptic depression at PV–PYR synapses through AP narrowing (22, 49). We demonstrate that a silent ion channel subunit engages these dynamics to modulate PV inhibition in a firing-frequency-dependent manner, a mechanism that is particularly relevant to fast-spiking interneurons.

The observed increase in paired-pulse ratios in *Kcng4*-knockout neurons is consistent with a reduction in release probability per release site. The loss of Kv6.4 shortens spike duration measured at the soma (**Fig. 3**, **Fig. S5**). If this effect is also present in the axon, it likely modifies presynaptic Ca^2+^ entry by decreasing the number of voltage-gated calcium channels opened per AP, thus altering both release probability and paired-pulse ratios. However, the peak amplitude of IPSCs in PYR neurons evoked by a single AP in a neighboring PV neuron was unaffected by loss of Kv6.4, suggesting potential compensatory changes in the number of release sites or postsynaptic GABA receptors per PV input. Nevertheless, our results do not establish the mechanisms by which Kv6.4 alters synaptic transmission between PV and PYR neurons. Additional direct or compensatory effects of Kv6.4 loss on Ca^2+^ entry, Ca^2+^ handling, or the release apparatus may also contribute. Future experiments directly measuring presynaptic Ca^2+^ entry can help discriminate between these mechanisms.

Kv6.4 is expected to share Kv2 subcellular localization patterns, forming clusters at the somatodendritic membrane (43, 44). The recent availability of effective Kv6.4 antibodies (36) will enable direct assessment of its localization in PV neurons. The broader expression pattern of Kv2.1 supports regulation of excitability across cell types, whereas the selective enrichment of Kv6.4 in PV neurons enables precise, cell-type-specific control of inhibitory output during high-frequency firing.

### *Kcng4*+ as a functional PV subclass in cortical development and plasticity

We find that the Kv6.4 silent subunit is selectively enriched in mature PV neurons—with minimal expression in other cortical cell types aside from astrocytes—and is restricted to a distinct transcriptional supertype of PV neurons (10). This specificity distinguishes Kv6.4 from broadly expressed potassium channels, such as Kv2.1 or Kv3.1 (3), making it an attractive candidate for selective targeting of this PV subclass. *Kcng4* has been identified as a marker of a PV subclass in the GPe (37), where *Kcng4*+ neurons were implicated in motor pattern regulation, but their physiological properties had not been defined. Here, we link this molecularly defined PV supertype to distinct physiological properties. Recent efforts in the molecular targeting of inhibitory neuron types have advanced our ability to dissect and manipulate circuit function across species (50). Kv6.4 thus represents a molecular marker of a specialized PV subclass with previously uncharacterized physiological roles, offering a new axis for selective circuit interrogation and tool development.

Kv6.4 expression confers a frequency-dependent mechanism for modulating PV output, which may facilitate adaptive changes in cortical circuits given that PV neurons form dense, nonspecific wiring with PYR neurons (51). *Kcng4* expression broadly parallels the spatiotemporal pattern of *Pvalb* but is particularly enriched in S1 L4, where sensory inputs first reach somatosensory cortex (52) and where inhibition mediates experience-dependent plasticity during critical periods (17, 52). The onset of Kv6.4 expression may also influence early sensory-driven plasticity (52). Indeed, PV-mediated experience-dependent plasticity in S1 is regulated by potassium channel function (18) and transcription (26). The role of Kv6.4 may be particularly relevant to motor and sensory learning, as we and others have found that Kv6.4 is enriched in mouse motor and sensory areas (36) and is involved in sensing pain (32, 34, 35). Thus, Kv6.4 could link short-term synaptic plasticity with long-term learning, and disruptions in its function could interfere with cortical excitation-inhibition, plasticity, and development.

### Pathophysiological implications of Kv6.4 dysfunction

Cortical circuit dysfunction in developmental and neurodegenerative disorders often arises from dysregulation of ion channels and GABAergic inhibition. Several voltage-gated ion channels and their regulators have been implicated in autism spectrum disorder, epilepsy, intellectual disability, and Alzheimer’s disease, such as the voltage-gated sodium channels Nav1.1 and Nav1.2 (50, 53), the synaptic adhesion molecule Cntnap4 (54), the Kv3 channel family (55), and NRG1-ErbB4 signaling that modulates Kv1.1 channels (25). These mechanisms operate within defined developmental windows or neuronal populations, such as PV neurons, to regulate excitability, plasticity, and network stability, and the disruption of these functions can result in abnormal sensory processing and cognitive deficits.

Kv6.4 has recently emerged as a clinically relevant modulator of neuronal excitability, motivating further investigation into its role in PV neurons *in vivo*. Loss-of-function *Kcng4* mutations in humans have been associated with reduced pain sensitivity in migraine (28, 34, 35) and labor during pregnancy (32), likely due to altered function of Kv2.1/Kv6.4 heterotetramers and disrupted excitability. Kv2.1/Kv6.4 complexes exhibit reduced sensitivity to channel blockers compared with Kv2.1 alone (33), and new pharmacological tools can distinguish silent Kv subunits from standard delayed rectifiers (56). These approaches can be leveraged to identify and target Kv6.4-containing complexes for functional or therapeutic studies. Complete Kv6.4 loss is poorly tolerated, as homozygous mutations are associated with male sterility (57) and hydrocephalus (58) in mice, suggesting that dosage-sensitive or cell-type-specific approaches may be necessary for potential interventions. Mutations in Kv2 channels resulting in complete or partial channel loss-of-function are linked to epilepsy and neurodevelopmental disorders such as autism spectrum disorder and intellectual disability, marked by seizures, motor and language problems, behavioral deficits, and ataxia (28, 59–62). Homozygous *Kcnb1* deletion in mice results in hippocampal hyperexcitability, epileptiform activity, accelerated seizure progression, locomotor hyperactivity, and impaired spatial learning (63). Future studies could examine how Kv6.4/Kv2.1 heterotetramers govern the timing and strength of PV-mediated inhibition during sensory processing and learning *in vivo*, and how these mechanisms are disrupted in disease.

## Materials and Methods

### Mice

We used Kcng4^Cre^ (The Jackson Laboratory, RRID:IMSR_JAX:029414) and PV^Cre^ (The Jackson Laboratory, RRID:IMSR_JAX:017320) mouse lines in this study. Both lines were crossed to the Ai14 reporter line (The Jackson Laboratory, RRID:IMSR_JAX:007914) to fluorescently label cells with tdTomato following Cre-mediated recombination. All Ai14 reporter animals (B6.Cg background) were heterozygous for the Ai14 allele. Kcng4^Cre^ (B6.129(SJL) background) is a knock-in/knock-out line in which Cre recombinase is inserted into exon 2 of the *Kcng4* gene, disrupting endogenous *Kcng4* gene expression. Kcng4^Cre^ animals used in this study were heterozygous (Kcng4^Cre/WT^) or homozygous (Kcng4^Cre/Cre^) for the Cre allele in addition to carrying one allele of Ai14. PV^Cre^ (B6.129P2 background) is a knock-in line in which Cre recombinase is inserted downstream of the *Pvalb* gene, without disrupting endogenous *Pvalb* gene expression. PV^Cre^ animals used in this study were heterozygous for the Cre allele (PV^Cre/WT^) in addition to carrying one allele of Ai14. These animals, which retain two copies of *Kcng4*, were used as wildtype control mice with labelling of PV neurons for experiments that require PV neuron identification (e.g. whole-cell patch-clamp recordings). Additionally, we used C57BL/6J mice (The Jackson Laboratory, RRID:IMSR_JAX:000664) as wildtype control mice and for backcrossing.

We used mice between 2 and 20 weeks of age, with approximately balanced sex ratios. Mice were maintained on a 12h light/dark cycle under standard housing conditions. All experiments were performed in accordance with protocols approved by the Harvard Standing Committee on Animal Care (Protocol #: IS00000571-6), following guidelines described in the US NIH Guide for the Care and Use of Laboratory Animals.

### Fluorescent RNA in situ hybridization

Wildtype mice (5–7 weeks old) were anesthetized with isoflurane, and brains were rapidly extracted and embedded in freezing media (Tissue-Tek O.C.T. Compound) on dry ice. Fresh-frozen brains were cryosectioned at 20 μm thickness (Leica CM3050S), adhered to SuperFrost Plus glass slides (VWR), and immediately refrozen, with short-term sample storage at –20°C and long-term sample storage at –80°C. Multiplexed fluorescent RNA in situ hybridization (FISH) was performed using the RNAscope Reagent Kit v2 (Advanced Cell Diagnostics, ACD) in collaboration with the Harvard Medical School Neurobiology Imaging Facility. The tissue was processed according to manufacturer instructions using a Protease III treatment for 30 min at room temperature. The following probes were used: Mm-*Pvalb* (ACD 421931), Mm-*Kcng4* (ACD 316931-C2), and Mm-*Kcnb1* (ACD 316941-C3). Samples were validated using positive and negative control probes, and nuclei were counterstained with DAPI. Samples were then imaged on an Olympus VS200 slide scanning microscope using a 20x objective and a Leica Stellaris 8 confocal microscope with a 63x oil-immersion objective. ROIs were drawn manually and quantified using QuPath, with further analysis in MATLAB (R2024a).

### Histology

Mice (2-20 weeks old) were anesthetized with isoflurane and transcardially perfused with 4% paraformaldehyde (PFA) in phosphate-buffered saline (PBS). Brains were dissected and postfixed in 4% PFA overnight at 4°C and transferred to 30% sucrose for at least 1 day prior to cryosectioning. Brains were embedded in freezing media (Tissue-Tek O.C.T. Compound), coronally sectioned at 50 μm thickness (Leica CM3050S), and stored in PBS at 4°C. Sections were stained with DAPI (Sigma-Aldrich D9542) for 10 minutes at room temperature, mounted on SuperFrost Plus glass slides (VWR), and coverslipped with mounting medium (ProLong™ Gold Antifade Mountant). Samples were imaged on an Olympus VS200 slide scanning microscope using a 20x objective and a Leica Stellaris 8 confocal microscope with a 63x oil-immersion objective.

### Acute slice preparation

Mice (5-7 weeks old) were anesthetized with isoflurane and immediately decapitated following transcardial perfusion with ice-cold artificial cerebrospinal fluid (ACSF) containing (in mM): 125 NaCl, 2.5 KCl, 25 NaHCO_3_, 2 CaCl_2_, 1 MgCl_2_, 1.25 NaH_2_PO_4_, and 17 glucose (300–310 mOsm kg^−1^). Extracted brains were transferred to a Leica VT1200s vibratome for preparation of 300 μm coronal slices in ice-cold ACSF. After cutting, slices were incubated for 3–6 min in choline-based solution at 34°C containing (in mM): 110 choline chloride, 25 NaHCO_3_, 2.5 KCl, 7 MgCl_2_, 0.5 CaCl_2_, 1.25 NaH_2_PO_4_, 25 glucose, 11.6 ascorbic acid, and 3.1 pyruvic acid (320–330 mOsm kg^−1^). In a second chamber, slices were incubated in ACSF at 34°C for 1 h. The slices were then cooled to room temperature (> 1h) and maintained until electrophysiology recordings were performed. All solutions were continuously bubbled with carbogen gas (95% O_2_, 5% CO_2_).

Whole-cell patch-clamp recordings were conducted in L2/3 of primary somatosensory cortex (S1) in artificial cerebrospinal fluid (ACSF) at 30–32°C with a perfusion rate of 4–6 mL min^−1^. Layer 2/3 was defined as spanning from 100–200 µm to 400–500 µm below pia. tdTomato+ neurons were identified using a Scientifica microscope equipped with a digital camera (Hamamatsu), an epifluorescence LED light source (CoolLED), and a 60x/0.9NA LUMPlanFl/IR water-immersion objective (Olympus). Patch pipettes were pulled from borosilicate glass (Sutter Instruments) with resistances of 2.0–3.5 MΩ. Recordings were performed using a Multiclamp 700B amplifier (Molecular Devices) and a data acquisition board (National Instruments). Data were collected and analyzed using MATLAB custom-written software and scripts (R2023a).

### Intrinsic properties

For whole-cell current-clamp recordings assessing intrinsic properties, patch pipettes were filled with low chloride potassium-based internal solution (in mM): 135 KMeSO_3_, 3 KCl, 10 HEPES, 1 EGTA, 0.1 CaCl_2_, 4 Mg-ATP, 0.3 Na-GTP, and 8 Na_2_Phosphocreatine (pH 7.3 adjusted with KOH, 290–300 mOsm kg^−1^). The following synaptic blockers were bath-applied: NBQX (10 μM, Tocris), CPP (10 μM, Tocris), and SR 95531 hydrobromide (10 μM, Tocris). Signals were filtered at 10 kHz and digitized at 50 kHz.

The experimenter was blinded to heterozygous (Kcng4^Cre/WT^; Ai14) and homozygous (Kcng4^Cre/Cre^; Ai14) genotypes, and age-matched littermates were used. Neuronal responses were measured to current steps delivered at both (1) the neuron’s endogenous resting membrane potential and (2) at –70 mV maintained via continuous current injection. Resting membrane potential was measured shortly after breaking in, after which all other intrinsic properties were measured. Action potential threshold was defined as the membrane potential at which the dV/dt exceeded 4% of the maximum slope during the upstroke.

No series resistance compensation was used. Series resistance did not exceed 20 MΩ. As provided in **Dataset S2**, mean resting membrane potential was –74.65 ± 1.02 mV, mean series resistance was 11.11 ± 0.40 MΩ, and mean membrane resistance was 71.14 ± 5.94 MΩ for PV^Cre/WT^ animals; –74.09 ± 1.36 mV, 11.18 ± 0.56 MΩ, and 67.37 ± 5.86 MΩ for Kcng4^Cre/WT^ animals; and –73.55 ± 0.88 mV, 10.71 ± 0.36 MΩ, and 63.94 ± 5.78 MΩ for Kcng4^Cre/Cre^ animals.

### Synaptic properties

For whole-cell voltage-clamp recordings assessing synaptic properties, patch pipettes were filled with high chloride cesium-based internal solution (in mM): 125 CsCl, 10 TEA-Cl, 10 HEPES, 0.1 EGTA, 3.3 QX-314, 1.8 MgCl_2_, 4 Mg-ATP, 0.3 Na-GTP, and 8 Na_2_Phosphocreatine (pH 7.3 adjusted with CsOH, 290–300 mOsm kg^−1^). The following synaptic blockers were bath-applied: NBQX (10 μM, Tocris) and CPP (10 μM, Tocris). Signals were filtered at 10 kHz and digitized at 10 kHz.

Cells in PV–PYR pairs were typically < 50 μm apart. Action potentials were evoked in PV neurons at the minimal current injection required between 1–2 nA (1 ms) to elicit a postsynaptic current in the PYR neuron. Absolute amplitude was measured shortly after breaking in via 3–5 single action potential stimulation pulses. Afterwards, eleven distinct action potential stimulation patterns were applied, each of which was repeated 3–5 times per paired recording in a randomized order. For analysis of paired-pulse recordings, overlap between the first and second IPSCs was corrected by subtracting the residual decay of the first IPSC from the peak of the second IPSC. No correction was applied to 5-pulse trains.

Again, no series resistance compensation was used during the experiment. Series resistance did not exceed 30 MΩ. For all paired recordings, holding current did not exceed 200 pA at −70 mV. Mean series resistance was 18.93 ± 0.90 MΩ and mean membrane resistance was 210.90 ± 20.50 MΩ for PYR neurons recorded from PV^Cre/WT^ animals; mean series resistance was 20.40 ± 1.51 MΩ and mean membrane resistance was 204.11 ± 11.80 MΩ for PYR neurons recorded from Kcng4^Cre/Cre^ animals.

IPSC peak amplitude was converted to conductance according to established procedures to compensate for variability in parameters such as series resistance and membrane resistance (64). Given that we are only measuring inhibitory current gi(t) and there is no excitatory current ge(t):

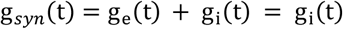

We used the following three equations to solve for gi(t):

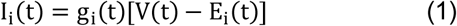

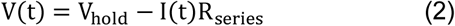

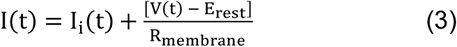

We derived the following equation for gi(t):

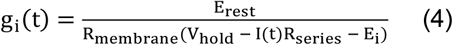

We measured Rmembrane and Rseries from a test pulse during the recording, and we separately measured Erest (–16 mV) and Ei (–2.5 mV). The PYR neuron was held at –70 mV; thus Vhold = –70 mV. Using these parameters, we converted inhibitory current I(t) to conductance gi(t).

### Quantification and statistical analysis

Data is plotted as mean ± SEM. P values are represented as * for P < 0.05 and ** P < 0.005, and details regarding the *p* values and statistical tests are provided in the figure legends and supplementary files.

### Data, materials, and software availability

All study data are included in the article and/or SI Appendix. The original code used to analyze the data in this study is publicly available on GitHub: https://github.com/sanikaganesh/IntrinSynC. Any additional information regarding the data and analysis is available from the lead contact upon request.

## Acknowledgments

We thank all members of the Sabatini lab for helpful advice and technical assistance. We are grateful to W. Wang for advice on electrophysiology experiments. We thank C. King for mouse colony management and assistance with blinding the experimenter to mouse genotypes. We also thank C. Chen, W. Regehr, G. Fishell, and M. Andermann for useful insights. Additionally, we thank M. Ocana, M. EL-Rifai, and the Neurobiology Imaging Facility at Harvard Medical School for collaborating with our lab to complete the FISH experiments. We are especially grateful to M. Sherman, J. Wallace, W. Wang, T. Kula, J. McMahon, S. Liu, A. Girasole, and K. Mastro for feedback on the manuscript.

This work was supported by NIH R35NS137336 (to B.L.S), National Science Foundation Graduate Research Fellowship Program (NSF–GRFP) DGE 2140743 (to S.G.) and the Herchel Smith Graduate Fellowship (to S.G.).

## Author Contributions

S.G. and B.L.S. designed research; S.G. and T.C. performed research; S.G., T.C., and B.L.S. analyzed data; S.G. and B.L.S. wrote the paper.

---

**Fig. S1.**
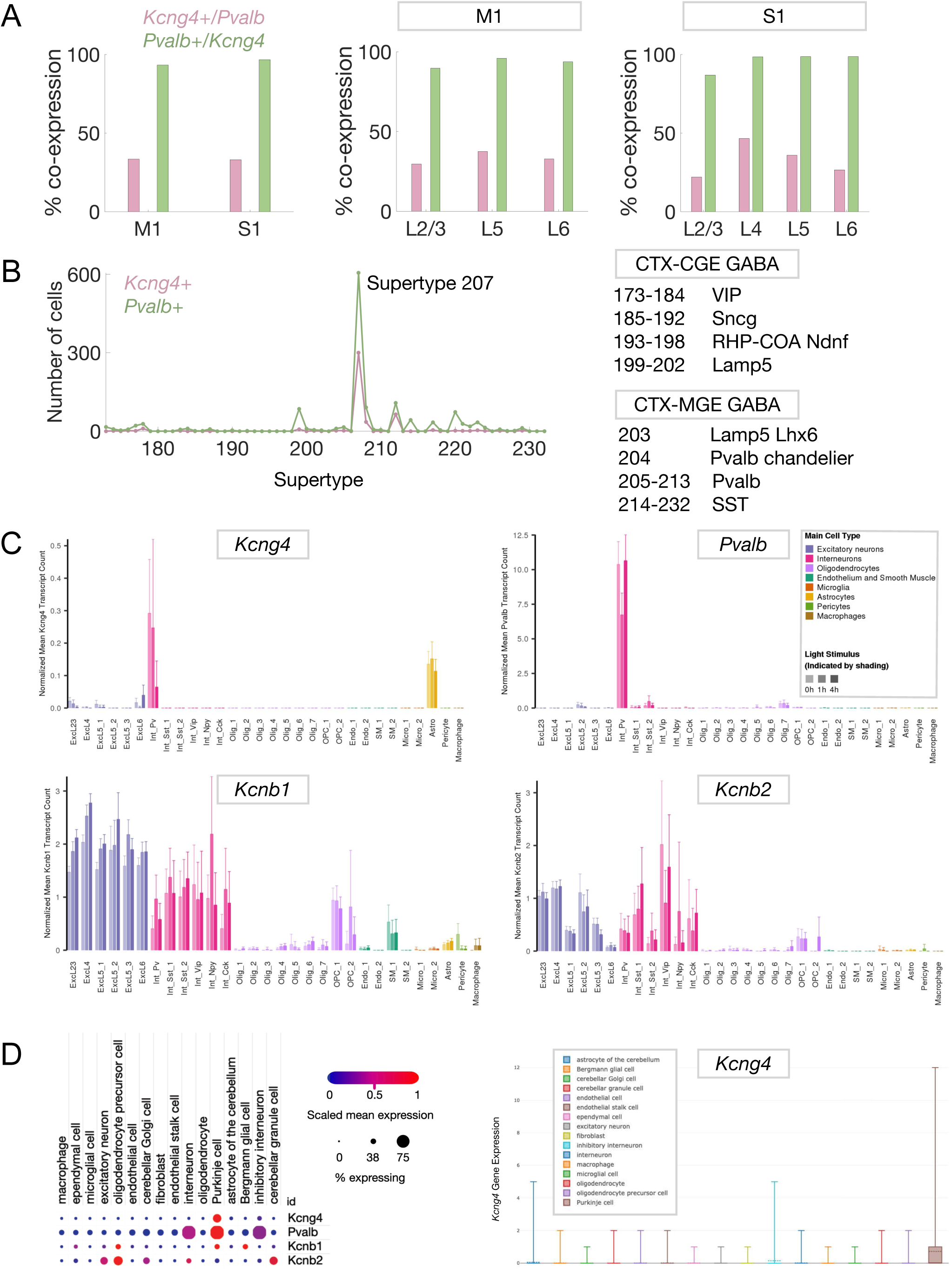
Co-expression and cell-type-specificity of *Kcng4* and *Pvalb* across spatial and single-cell transcriptomic datasets. **A**, Quantification of cells co-expressing *Kcng4* and *Pvalb* in adult mouse cortex was performed using Multiplexed Error-Robust Fluorescence *In Situ* Hybridization (MERFISH) data from the Allen Brain Cell (ABC) Atlas (Zhuang-ABCA-1, MERFISH whole brain coronal 1), accessed via the Brain Knowledge Platform (10). Percentage of co-expressing cells in M1 and S1, labeling as in **Fig. 1E**. **B**, Number of *Kcng4*- and *Pvalb*-expressing cells in S1 across PV neuron supertypes, as classified by the ABC Atlas (*left*). Legend detailing inhibitory subtypes (*right*). Abbreviations: CTX, cortex; CGE, caudal ganglionic eminence; MGE, medial ganglionic eminence; GABA, GABAergic neuron; VIP, vasoactive intestinal peptide; Sncg, synuclein, gamma; RHP, retrohippocampal region; COA, cortical amygdalar area; Ndnf, neuron-derived neurotrophic factor; Lamp5, lysosomal-associated membrane protein family, member 5; Lhx6, LIM homeobox protein 6; Pvalb, parvalbumin; SST, somatostatin. **C**, Normalized transcript counts for *Kcng4*, *Pvalb*, *Kcnb1*, and *Kcnb2* across cell types in mouse adult visual cortex from single-cell RNA sequencing data (3) accessed via an interactive gene expression database. **D**, Gene expression for *Kcng4*, *Pvalb*, *Kcnb1*, and *Kcnb2* across cell types in mouse adult cerebellar cortex from single-nuclei RNA sequencing data (6) with a detailed visualization of *Kcng4* gene expression, accessed via the interactive Single Cell Portal.

**Fig. S2.**
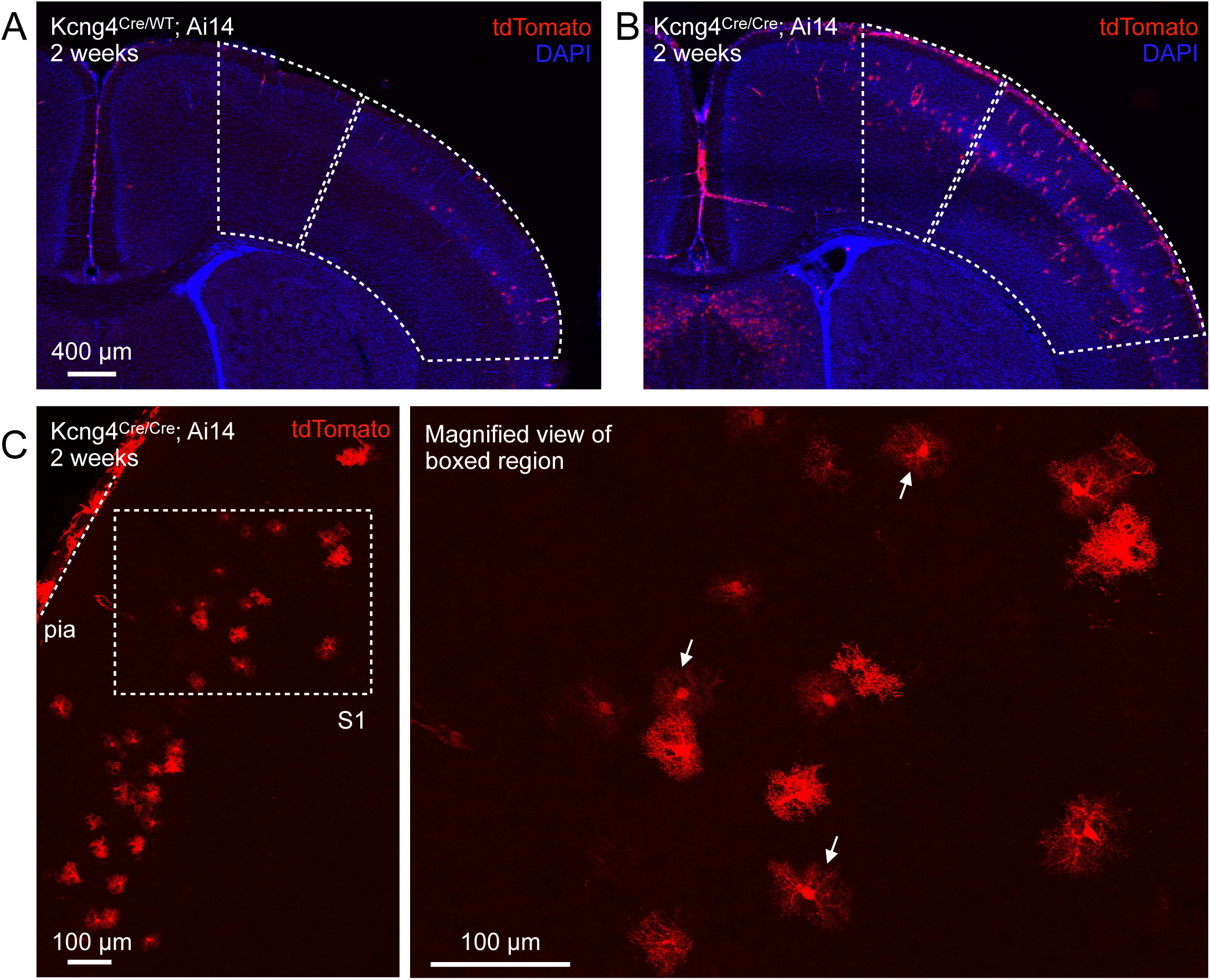
Kcng4^Cre^ activation in juvenile mice. **A-B**, Representative images showing tdTomato fluorescence in M1 and S1 for heterozygous (Kcng4^Cre/WT^; Ai14) **(A)** and homozygous (Kcng4^Cre/Cre^; Ai14) **(B)** mice at 2 weeks, corresponding to the regions outlined in **Fig. 2A**. **C**, Higher magnification images illustrating the distribution of tdTomato-positive cells in S1 of a homozygous (Kcng4^Cre/Cre^; Ai14) mouse. Magnified view (*right*) of the boxed region (*left*) marked by dashed white lines. Solid white arrows highlight example cells exhibiting astrocytic morphology (*right*).

**Fig. S3.**
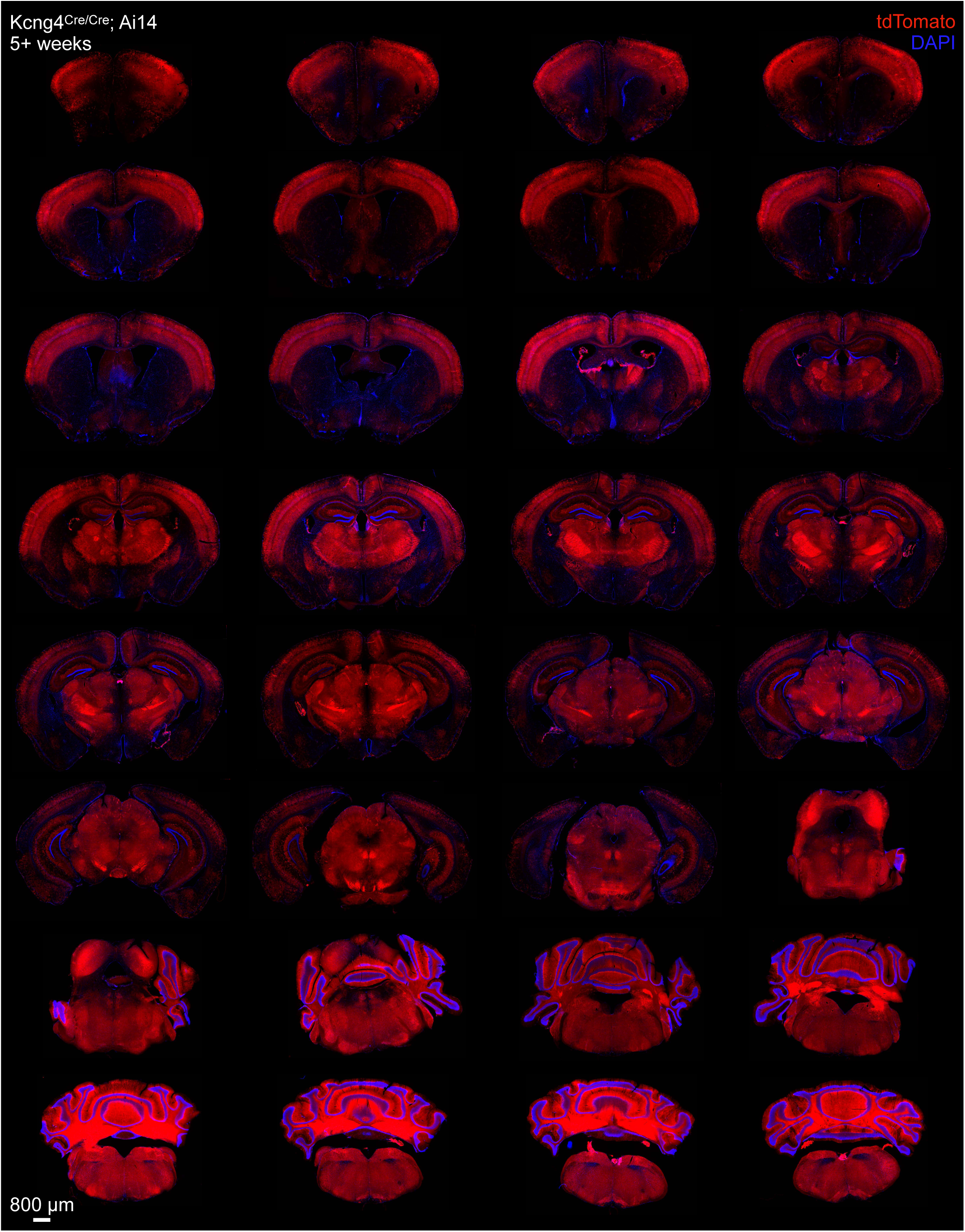
Brain-wide Kcng4^Cre^ activation in adult mice. Representative images showing tdTomato fluorescence in a homozygous (Kcng4^Cre/Cre^; Ai14) mouse at 5+ weeks across various brain areas, arranged anterior to posterior.

**Fig. S4.**
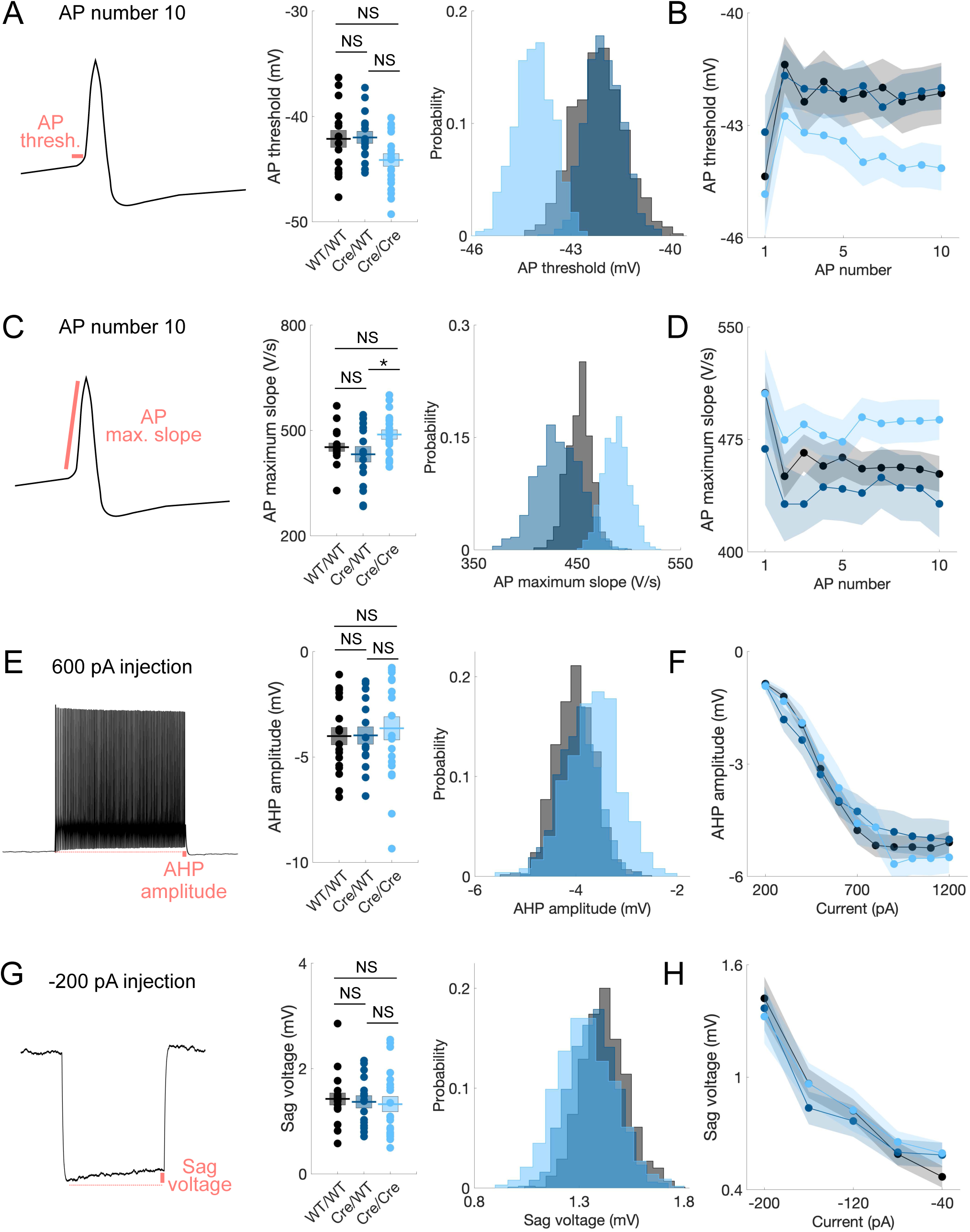
Additional intrinsic properties of PV neurons following Kv6.4 deletion. **A**, Schematic of AP threshold (*left*), group means (mV, *center*), and bootstrapped distribution of the means (*right*) for the 10^th^ AP, as described in **Fig. 3**. **B**, Mean values across the first 10 APs (mV). **C-H**, As in **A** and **B** for AP maximum slope (V/s) (**C-D**), afterhyperpolarization (AHP) amplitude (mV) (**E-F**), and sag voltage (mV) (**G-H**). AHP amplitude was measured from a current injection of 600 pA (**E**) and across stepwise current injections from 200 to 1200 pA (**F**). AHP amplitude was defined as the difference between the resting membrane potential and the most negative voltage following current injection. Sag voltage was measured from a current injection of –200 pA (**G**) and across hyperpolarizing steps from – 200 to –40 pA (**H**). Sag voltage was defined as the difference between the steady-state voltage and peak hyperpolarization during hyperpolarizing current injections. Mean ± SEM displayed. * *P* < 0.05; NS: not statistically significant. One-way ANOVA (Tukey-Kramer post hoc multiple comparisons). Bootstrapping (1,000 iterations) was used to visualize sample mean distributions.

**Fig. S5.**
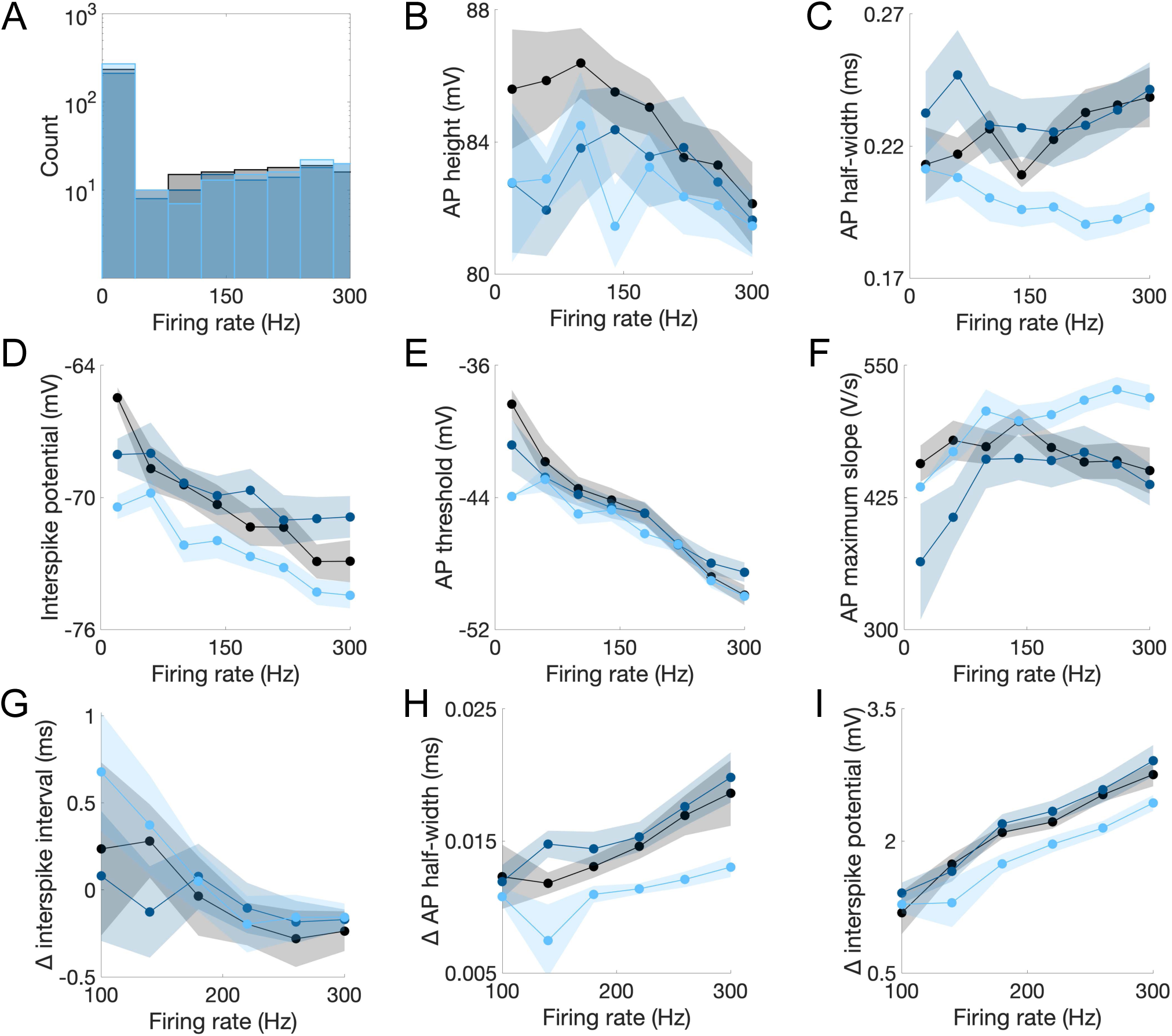
Extended analysis of PV intrinsic properties following Kv6.4 deletion. **A**, Histogram of firing rate (Hz) distribution (within 300 Hz) for all recordings, independent of current injection, across groups described in **Fig. 3**. **B**, AP height (mV) across firing rate for all recordings. **C-F**, As in **B** for AP half-width (ms) (**C**), interspike potential (mV) (**D**), AP threshold (mV) (**E**), and AP maximum slope (V/s) (**F**). **G**, Change (Δ) in interspike interval (ISI; ms), defined as ISI49-50 – ISI2-3, across firing rate in recordings containing ≥50 APs. **H-I**, As in **G**, for Δ AP half-width (ms), defined as AP50 half-width – AP2 half-width (**H**), and Δ interspike potential (mV), defined as AP50 interspike potential – AP2 interspike potential (**I**). Mean ± SEM displayed.

**Fig. S6.**
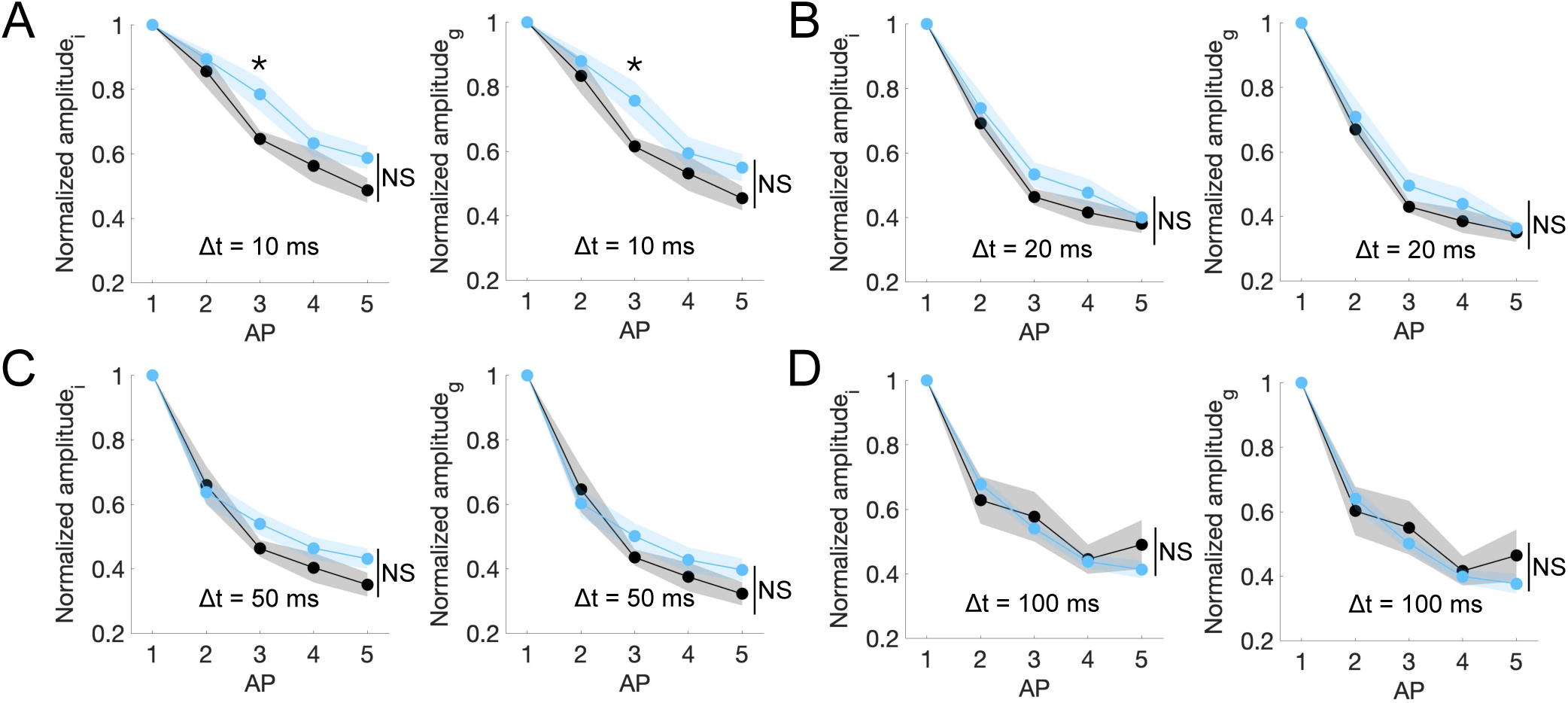
Paired pulse depression at PV-PYR synapses across varying interstimulus intervals following Kv6.4 deletion. **A-D**, Amplitude of inhibitory current (i; *left*) and conductance (g; *right*) across groups, normalized to response from the first AP, during trains of five presynaptic action potentials, as described in **Fig. 4**. Trains were delivered with an ISI (Δt) of 10 ms (**A**), 20 ms (**B**), 50 ms (**C**), and 100 ms (**D**). Mean ± SEM displayed. * *P* < 0.05; NS: not statistically significant. Unpaired two-tailed t-tests or repeated-measures ANOVA (Greenhouse-Geisser corrected).

**Datasets**

Dataset S1 (separate file). Raw data and summary data for Fig. 1 and Fig. S1.

Dataset S2 (separate file). Summary data for Fig. 3 and Fig. S4.

Dataset S3 (separate file). Summary data for Fig. 4 and Fig. S6.

